# Successes of artemisinin elicitation in low-artemisinin producing *Artemisia annua* cell cultures constrained by repression of biosynthetic genes

**DOI:** 10.1101/740167

**Authors:** Melissa Kam Yit Yee, Winnie Yap Soo Ping

## Abstract

The sesquiterpene phytolactone derived from *Artemisia annua*, artemisinin is associated with a variety of novel biological properties, such as immunoregulatory and anticancer effects, and therapeutic applications, apart from its main function as an antimalarial drug. Emerging from the fact that artemisinin production *in planta* occurs in trace amounts and its compartmentalized synthesis, the irregular agricultural supply often results in market fluctuations and reductions in artemisinin inventory. Further improvement in artemisinin production calls for approaches that act in a supplementary manner, filling the agricultural production gap. Here we investigated the elicitation efficiency of ultraviolet B (UV-B) and dimethyl sulfoxide (DMSO) independently on a low-artemisinin producing (LAP) chemotype of the species *A. annua*. The exposure of cell suspension cultures to short-term UV-B radiation and DMSO treatment did not result in significant changes in artemisinin yield. The lack of stimulation has been associated with: (i) the general lack of cytodifferentiation of cell cultures; (ii) negative feedback regulation of artemisinin biosynthesis; and (iii) artemisinin sequestration by cellular detoxification. Further molecular analysis revealed the repression of key genes *ADS, DBR2* and *ALDH1* which affected artemisinin synthesis. This study provides insights into the complexity of stress-induced responses of *A. annua* cell suspension cultures in relation to metabolic processes (transportation, accumulation and degradation of secondary products) which are important for artemisinin formation.

## Introduction

The highly oxygenated endoperoxide sesquiterpene lactone, artemisinin is an effective antimalarial compound that is derived from the Chinese medicinal plant *Artemisia annua* L. In view of the rapid development of parasite resistance to former antimalarial drugs, the World Health Organization (WHO) has endorsed the artemisinin-based combination therapies (ACTs) as the first-line treatment for uncomplicated malaria caused by *Plasmodium falciparum* [1, 2]. Apart from its antimalarial activities, artemisinin is also a multi-functional compound that is associated with a variety of novel biological properties, such as immunoregulatory [3], anticancer [4, 5], antituberculosis [6], antidiabetic [7], antiparasitic and antiviral functions [8].

Given the economic importance of artemisinin, the sole dependency on plant-based production periodically fails to meet the global demand due to irregular agricultural supply, which then renders fluctuations in the market and reductions in artemisinin inventory [9, 10]. To mitigate supply uncertainties and price volatility, biotechnological alternatives such as genetic modification, crop breeding and semi-synthetic production were considered with the latter far more successful [11]. The semi-synthesis approach has the production capacity to yield approximately 60 tonnes of artemisinin annually, but it also involves the trade-off between efficiency and profitability. The multiple chemical conversion steps to produce a semi-synthetic variant of artemisinin, compared to the more economical leaf-derived artemisinin extraction, demands high financial and environmental costs, inevitably increasing raw material prices [12, 13]. Furthermore, demand has plateaued over the last two years due to excess agricultural artemisinin and improved malaria diagnosis, plant-derived artemisinin was sold at a price less than US$250 per kilogram, while semi-synthetic artemisinin was valued between US$350 – 400 per kilogram [14]. Nor was it a cheaper alternative to the existing agricultural source. Semi-synthetic artemisinin is neither alleviating shortages. As sufficient supply of raw materials can be produced by farmers in good years, an economic complementary production strategy which is also rapid in establishment should be considered to address the agricultural production gap.

Naturally, artemisinin biosynthesis occurs in glandular secretory trichomes that are present on foliage, stems and inflorescences of the *A. annua* plant [15, 16]. Due to its compartmentalized synthesis and influence by physiological, seasonal and environmental factors, artemisinin concentration is highly variable with reported values of 0.1 – 10 mg g^-1^ dry weight (DW) [11, 17]. With the biosynthetic pathway of artemisinin formation fully elucidated over the past decade, it was learnt that within the species *A. annua* two contrasting chemotypes can be distinguished, which are characterized by the differing contents of artemisinin and its precursor (Fig 1). While both possess artemisinin, the high-artemisinin producing (HAP) chemotype contains relatively higher levels of dihydroartemisinic acid (DHAA) and artemisinin, and the low-artemisinin producing (LAP) chemotype has higher contents of artemisinic acid and arteannuin B (a non-antimalarial product) [18, 19]. Despite the difference in biochemical phenotype which is attributed to the differential expression of one artemisinin biosynthesis specific gene artemisinic aldehyde Δ11(13) reductase (DBR2), both chemotypes undergo similar spontaneous photooxidation reactions to produce artemisinin or arteannuin B from its direct precursors [20–23].

**Fig 1.**
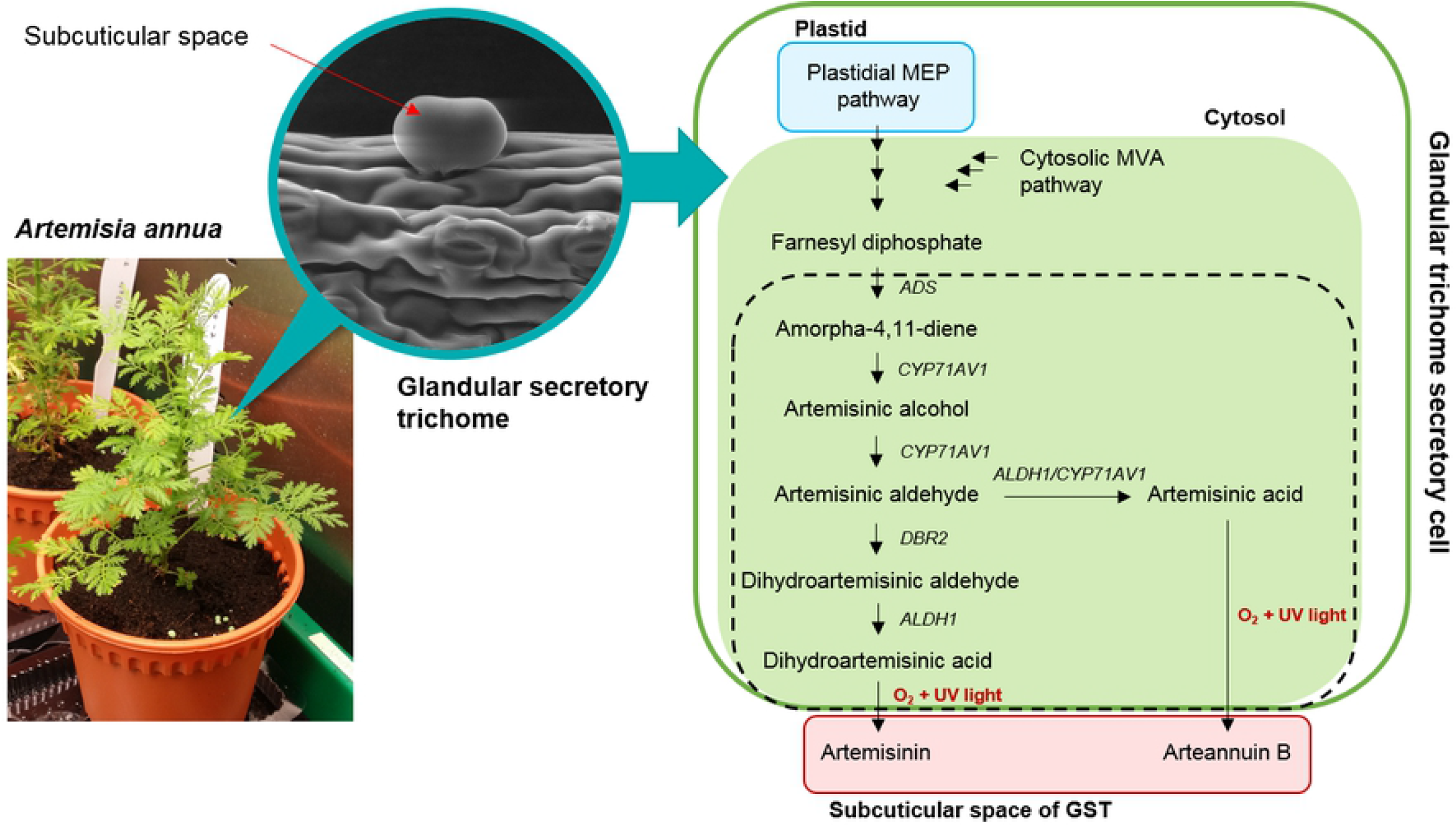
Schematic representation of artemisinin biosynthesis. Synthesis occurs in the glandular trichome secretory cells of *A. annua* and sequestration of artemisinin takes place in the subcuticular space. The two branching pathways leading to artemisinin and arteannuin B of the artemisinin biosynthetic pathway (dashed line box) are depicted and the final non-enzymatic photo-oxidation reaction is shown in red. MEP, methylerythritol 4-phosphate; MVA, mevalonate; GST, glandular secretory trichome; ADS, amorpha-4,11-diene synthase; CYP71AV1, cytochrome P450 monooxygenase; DBR2, artemisinic aldehyde Δ11(13) reductase; ALDH1, aldehyde dehydrogenase 1.

As a secondary metabolite used mainly in defence mechanisms, production of artemisinin would naturally be induced and enhanced under environmental stresses. Under such adverse conditions the elevation of secondary oxidative stress, which is a consequence of primary biotic/abiotic stress, is in a direct synergistic relationship with artemisinin production [24–27]. On such basis, the biotechnological method elicitation which mimics the natural induction of plant stress is the method of choice to enhance the production of artemisinin. The positive effects of UV-B and DMSO on the production of artemisinin had been verified by previous studies on *A. annua* seedlings [28, 29], *in vitro* propagated plantlets [30] and shoot cultures [31]. Former studies had also shown that artemisinin production can be enhanced by different elicitors in *A. annua* cell cultures [32–35]. However, to our knowledge, the effect of UV-B and DMSO elicitation on the contents of artemisinin synthesized by a LAP chemotype cell culture has not previously been reported. As LAP chemotype preferentially converts its precursor artemisinic acid to arteannuin B than artemisinin under normal circumstances, it seemed possible that stress factors could induce a shift in its biosynthetic mechanism favouring artemisinin formation. This notion is motivated by preliminary studies of the same stress treatments, but on young soil-grown LAP plants (S1 Table). Given that DHAA is a direct precursor of artemisinin with the implicated role as a ROS scavenger [36, 37], our hypothesis is that UV-B and DMSO applications on a LAP may trigger an oxidative burst in cells resulting in an increased DHAA production for ROS quenching, which in turn enhances the production of artemisinin. The main objective of the present study is therefore to investigate the above-mentioned hypothesis by evaluating the effects of UV-B and DMSO elicitation, respectively on the content of artemisinin in cell cultures of a LAP chemotype.

## Materials and methods

### Plant materials and culture conditions

Seeds of European origin obtained from Chiltern Seeds Limited (Oxfordshire, UK) were germinated and grown in sterile soil under controlled conditions. Seedlings were transplanted into pots once a transplantable size of approximately 15 cm was achieved. Plants were cultivated in a growth chamber with a regimen of 16 h day and 8 h night at 25°C to prolong the vegetative stage. The plants were watered every alternate day with direct application to the soil and fertilized once every 2 weeks. Stock plants were maintained and propagated via stem cuttings. Unsown seeds were stored at 4°C until further use.

For *in vitro-raised* stock plants, seeds were surface sterilized according to Pras et al. [38] with slight modifications. Seeds were disinfected in 3% (v/v) Clorox^®^ (commercial bleach containing 5.25% sodium hypochlorite) for 20 min with prior soaking in distilled water for 4 h. The seeds were rinsed three times with sterile distilled water and germinated on basal Murashige and Skoog (MS) medium [39] (Duchefa Biochemie B. V., Netherlands) containing 3% (w/v) sucrose (Fisher Scientific, USA) and 0.35% (w/v) Phytagel^™^ (Sigma-Aldrich, USA). Thereafter *in vitro*-raised plantlets were subcultured at 4-week interval into fresh basal MS medium with 3% (w/v) sucrose and 0.25% (w/v) Phytagel^™^. All media were adjusted to pH 5.8 and autoclaved at 121°C for 15 min. Cultures were incubated at 25 ± 2°C under a 16 h photoperiod (40 μmol m^-2^ s^-1^) provided by cool-white fluorescent lamps (Akari TLD36W/54).

### Callus induction

Young leaves of *in vitro*-raised plants (1- to 2-month-old) were excised (0.5 – 1.0 cm) and cultured onto MS medium supplemented with 0.5 mg L^-1^ 6-benzylaminopurine (BAP), 0.5 mg L^-1^ α-naphthaleneacetic acid (NAA) (Duchefa Biochemie B. V., Netherlands), 3% (w/v) sucrose and 0.35% (w/v) Phytagel^™^ [33]. After 3 weeks of callus initiation, friable calli were separated from explants and transferred to the same fresh medium for further proliferation. Subsequent maintenance of calli were performed at 4-week interval [40].

### Establishment of *A. annua* cell suspension

Fine cell suspension cultures were initiated with the same medium composition used for callus induction [41]. Friable calli (2 g) were inoculated into 100 mL Erlenmeyer flasks containing 25 mL liquid MS medium with 0.5 mg L^-1^ BAP and 0.5 mg L^-1^ NAA. Flasks were placed on an orbital shaker (120 rpm) and incubated under the same culture conditions as above. After 3 weeks of inoculation, established suspension cultures were filtered using a sterile stainless-steel sieve (aperture of 1.5 – 2.0 mm) to obtain a homogenous cell suspension. The fine cell suspension was then diluted with the same volume of fresh medium (dilution of 1:1 v/v), distributed into two 100 mL Erlenmeyer flasks and incubated in the same conditions. Medium volume was increased to 50 mL in the following subcultivation step. Homogenized suspension cultures (25 mL) were transferred into 250 mL Erlenmeyer flasks and refreshed by adding equal volumes of fresh medium (25 mL). Cell cultures were gently swirled and further incubated.

### Growth kinetics of suspension cultures

Growth of suspension cultures was evaluated by determining the cell dry weight during a culture cycle of 35 days with measurements taken at 5-day interval. Cells were harvested by filtration through filter paper (Sartorius^™^ Quantitative Grade 292 Filter Papers; Fisher Scientific, USA), rinsed with distilled water and blotted to remove excess water. The cells were then oven dried at 60°C for 24 h to determine the dry weight (DW) [42]. Triplicate flasks were used in this experiment.

### Cell viability analysis by MTT tetrazolium reduction assay

To evaluate viability of *A. annua* cells over the growth period of 35 days, the 3-(4,5-dimethylthiazol-2-yl)-2,5-diphenyltetrazolium bromide (MTT) tetrazolium reduction assay was carried out according to Castro-Concha et al. [43] with minor modifications. Suspension cells (1 mL) were washed twice with 50 mM phosphate buffer (pH 7.5) and resuspended in 1 mL of the same buffer. MTT (115 μL) was added to a final concentration of 1.25 mM. Samples were then incubated in darkness for 1 h at 37°C. To dissolve the formazan crystals, 1.5 mL solubilization solution (50% [v/v] MeOH containing 1% [w/v] sodium dodecyl sulfate) was added to the samples and incubated at 60°C for 30 min. Samples were centrifuged at 1880 ×g for 5 min at room temperature and supernatant recovered. Absorbance was quantified at 550 nm.

### UV-B elicitation and sample collection

The effects of UV-B treatment in enhancing the capacity of *A. annua* cell cultures to produce artemisinin were tested on 10-day-old suspension cultures (after subculture). Flasks were exposed to artificial UV-B radiation for 0, 1, 2, 3, 4 and 5 h, respectively. UV-B radiation was artificially provided by UV-B Narrowband TL lamp (TL 20W/01 RS SLV/25) (Philips Lighting Holding B. V., Netherlands) at a given dosage of 2.9 W m^-2^. Cells were harvested immediately by filtration after every hour of UV-B treatment for biochemical and molecular analysis.

### DMSO treatment and sample collection

DMSO concentrations were manipulated to evaluate impact on artemisinin production in 5-day-old suspension cultures (after subculture). Old liquid medium was removed by filtering through a 100 μm nylon cell strainer (Falcon^®^; BD Biosciences, USA), the trapped cells were then returned to the flask and 50 mL fresh liquid media supplemented with the same hormone regime was added. Thereafter, filter-sterilized DMSO (Sigma-Aldrich, USA) was added at increasing concentrations (0, 0.1, 0.25, 0.5, 1.0, 2.0% [v/v]). Flasks were cultured under the same growth conditions and cells were harvested one-week post-treatment.

### Artemisinin extraction and determination

For crude extract preparation, treated and untreated cells were harvested and dried in the same manner as described above, and ground into fine powder using a pestle and mortar. Artemisinin was extracted as described by Xiang et al. [44] with minor modifications. Briefly, 500 mg of dry powder were extracted with 50 mL of petroleum ether (analytical grade; Merck, Germany) in an ultrasonic bath for 30 min, the extraction mixture was filtered, followed by a second extraction with 50 mL of petroleum ether and sonicated for 10 min. Filtrates from both extractions were pooled. Employing a rotary evaporator (Büchi Rotavapor R-200, Switzerland), filtrates were evaporated to dryness *in vacuo* at 50°C. Residue was re-dissolved in 5 mL methanol (HPLC grade; Merck, Germany) and stored in −20°C. Prior to HPLC analysis, 1 mL of the solution was filtered through a 0.45 μm nylon membrane filter (Merck Millipore, USA).

Artemisinin detection was performed using an Agilent 1260 Infinity LC system equipped with a Hypersil GOLD C18, 250 x 4.6 mm column (pore size 175 Å, particle size 5 μm) (Thermo Scientific, USA) coupled with a guard column. The mobile phase consisted of acetonitrile (HPLC grade; Merck, Germany) and water (60:40 v/v) at a constant flow rate of 1 mL min^-1^. Injection volume and detection wavelength were set at 20 μL and 195 nm, respectively. Artemisinin standard (≥98%; LKT Laboratories Inc., USA), dissolved in methanol and properly diluted, was used to prepare standard curves for quantification. Putative artemisinin peaks were confirmed by re-analysing a sample co-injected with standard artemisinin. For each sample, three replications (n ≥ 3) were performed.

### Chemotype identification

Young leaves of soil-grown plants were used for genomic DNA extraction using innuPREP Plant DNA kit (Analytik Jena, Germany) following the manufacturer’s instruction. The 3’-end of the DBR2 promoter was amplified by PCR using DNA as template, and primers pDBR2_for_ and pDBR2_rev_ (S2 Table) [23]. The PCR system included 10 μM forward primer and 10 μM reverse primer, 0.5 μL of genomic DNA, 10 mM dNTP mix, 5 μL of 5× Green GoTaq^®^ Flexi Buffer and 0.25 μL of 5U GoTaq^®^ DNA Polymerase (Promega, USA) in a total volume of 25 μL. Amplification was conducted using Mastercycler^®^ nexus gradient (Eppendorf, Germany) under the following thermal cycling program: 3 min at 95°C; 30 cycles of 30 s at 95°C, 30 s at 65°C, 45 s at 72°C; 10 min at 72°C.

### RNA isolation, cDNA synthesis and PCR amplification

Pre-weighed cell samples of 100 mg were frozen in liquid nitrogen and ground to a fine powder using a sterile pestle and mortar. Total RNA was extracted using innuPREP Plant RNA kit (Analytik Jena, Germany). The purity and yield of RNA samples were measured with a NanoDrop™ 1000 (Thermo Scientific, USA) and integrity tested by 1% (w/v) agarose gel at 90V for 55 min. RNA samples were stored in −80°C until further use. First-strand cDNA was synthesized from 1 μg of total RNA using QuantiTect^®^ Reverse Transcription kit (Qiagen, Germany).

PCR amplification was performed using primers ADS_Ex4_for_ and ADS_Ex4_rev_ for the detection of DNA contamination, as the primers were designed to span intron 4 of the amorpha-4,11-diene synthase (*ADS*) gene. The amplification of a 251 bp fragment indicates the authenticity of the cDNA products that are contamination-free, whereas a single fragment of 363 bp indicates genomic DNA contamination [45]. Only samples that resulted in the amplification of the former were taken for further analysis. Routine detection of the artemisinin biosynthesis specific genes: ADS, *DBR2*, aldehyde dehydrogenase 1 (*ALDH1*), cytochrome P450 monooxygenase (*CYP71AV1*), cytochrome P450 reductase (*CPR*); and housekeeping genes: ubiquitin (*UBI*), actin 2 (*ACT2*) was carried out using Mastercycler^®^ nexus gradient. Reactions were performed in a total volume of 25 μL with 100 ng cDNA, 10 μM forward and reverse gene-specific primers and 5× Green GoTaq^®^ Flexi Buffer. Cycling conditions were 10 min at 95°C; 40 cycles of 30 s at 95°C, 30 s at 60°C, 30 s at 72°C. Amplified products (10 μL) were electrophoresed on 1% (w/v) agarose gel at 90V for 45 min. All primers are listed in S2 Table.

### Data analysis

Each of the two treatments (UV-B and DMSO) was replicated six and nine times, respectively with each flask treated as one replicate. Experiments were set up in a randomized complete block design. Significance of stress treatments was determined by Kruskal-Wallis test. P < 0.05 was considered to be statistically significant. All statistical analyses were performed using the SPSS software (IBM SPSS Statistics 24).

## Results and discussion

### Growth kinetics of *A. annua* cell suspension

The growth dynamics of *A. annua* cell suspension cultures were determined via dry cell weight estimation. It appeared that *A. annua* cells are better able to adapt to the new culture environment as evidenced by the rapid entry of cells into the exponential growth phase within the first 5 days of culture initiation (Fig 2A). After day 10 of culture, the cells then entered plateau, followed by a phase of diminishing growth. The inclusive of growth hormones (0.5 mg L^-1^ BAP and 0.5 mg L^-1^ NAA) in media for callus induction from leaf explants derived from *in vitro* plants cultured in basal MS, may have conditioned cells to better adapt and respond to the varying physicochemical conditions of its environment. Such phenomenon is known as priming in which a transient abiotic stress cue leads to modified stress responses upon exposure to a recurring stress [46]. Thus, following its transition into liquid suspension supplemented with the same hormonal regime, cells exhibit enhanced tolerance to recurring stress in terms of rapid growth as lag phase is often associated with minimal growth that is compromised for the exploitation of new environmental conditions [47, 48]. It is also likely that the lag phase has been limited to a duration of only 1–3 days when subculture is conducted during the exponential or linear phases [49]. The subculture period of 3-week interval of the present study may justify the shortened lag phase. Furthermore, Lo et al. reported prolonged lag phase (9 days) when Vietnamese *A. annua* suspension cultures with initial inoculum of 0.25 g were subcultured at a 2-week interval [50]. Based on the growth curve, exponentially growing *A. annua* cultures of day 5 and 10 were selected for DMSO and UV-B elicitation experiments, respectively.

**Fig 2.**
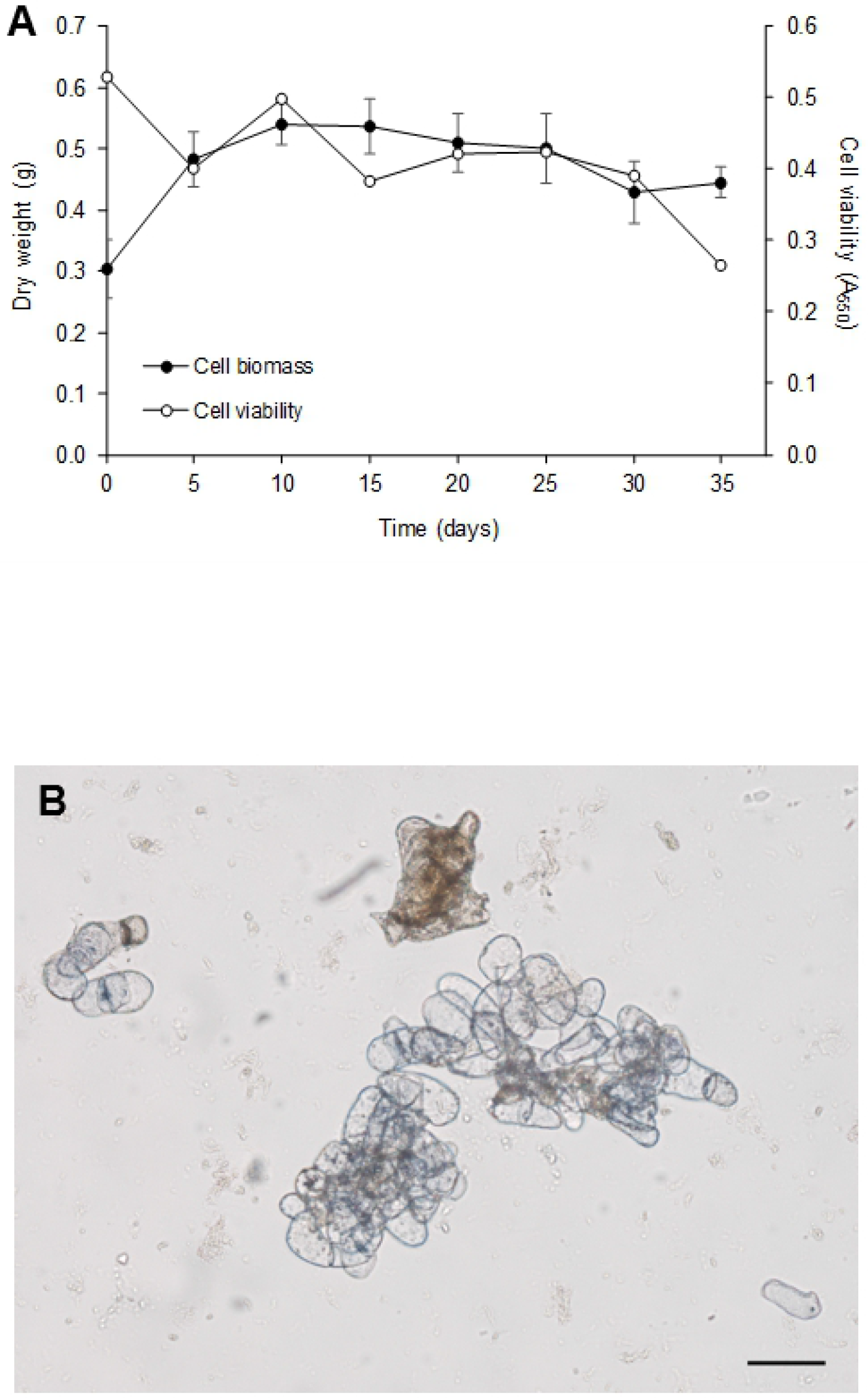
Evaluation of growth kinetics and MTT assay of *A. annua* suspension cultures. (A) Time profile of growth and viability of *A. annua* cell cultures in liquid MS medium supplemented with 0.5 mg L^-1^ BAP and 0.5 mg L^-1^ NAA in shake flasks. Values are mean ± standard deviation of triplicate cultures. (B) Viable single cell and cellular aggregates were distinguishable with purple cytoplasm, non-viable cells remained brown coloured. Scale bar represents 0.1 mm.

### Identification of chemotype

In the present investigation, the biochemical phenotype of the *A. annua* plant used in this study was validated using the DNA marker described by Yang et al. [23]. The selection method was developed based on the DBR2 promoter activity. The major difference between the chemotypes lies within their promoter sequences whereby low-artemisinin producers (LAPs) contain several deletions or insertions in the region upstream of the ATG start codon, while the rest of the sequences are highly similar to that of high-artemisinin producers (HAPs). Due to this promoter variation, expression levels of *DBR2* gene in LAP groups were compromised which in turn affects artemisinin biosynthesis. Therefore, amplification of the 3’-end of DBR2 promoter with specific primers would allow the discrimination between the two chemotypes. The results indicated that this unknown European variety belongs to the LAP group, concluded based on the absence of a 580 bp fragment characteristic of the HAP chemotype [23] (Fig 3). This is consistent with the general trend that LAP chemotype is mostly represented by *A. annua* varieties originating from Europe and North America [18]. It is reasonable to suggest the use of a LAP plant in our investigation as the direct effects of elicitors in enhancing artemisinin production can thus be accurately determined. Critical analysis of previous studies showed that chemotype of *A. annua* used are often not well informed, rendering direct comparison of results of similar experiments difficult. With the validation of its biochemical phenotype, the data obtained from this study can therefore be adopted into future research relating to LAPs.

**Fig 3.**
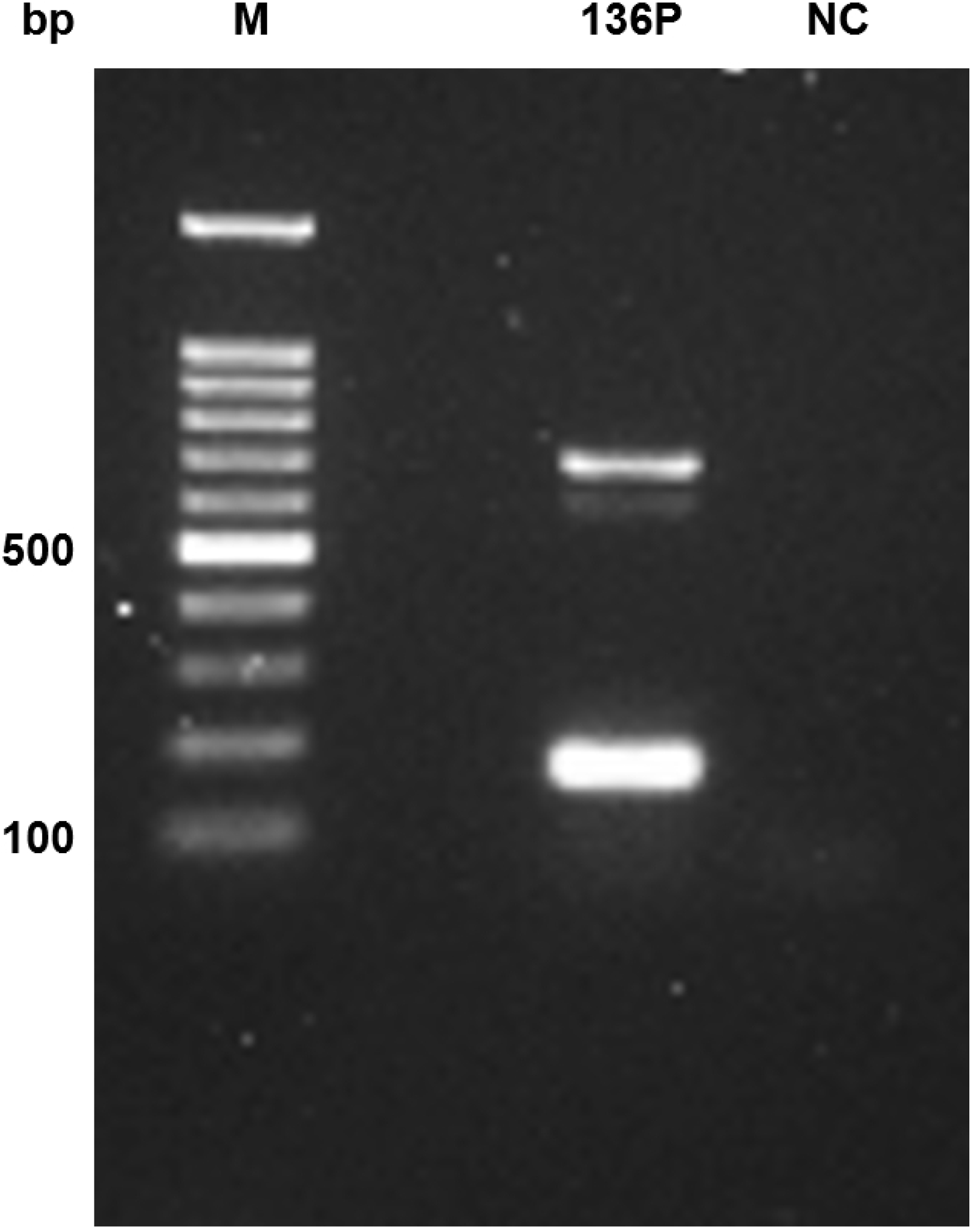
Agarose gel electrophoresis of amplified 3’-end of DBR2 promoter of *A. annua*. M, 100 bp DNA ladder; 136P, LAP *A. annua*; NC, negative control.

### Stress effects on the artemisinin content of *A. annua* suspension cells

To test our hypothesis that elicitation with physical and chemical stress may lead to a shift in the biosynthetic mechanism of a LAP favouring artemisinin formation instead of arteannuin B, *A. annua* suspension cultures were subjected to UV-B radiation and DMSO application independently. The effects of these stress treatments are shown in Table 1 and 2. In the present study, short term UV-B radiation showed only limited effect on the production of artemisinin with the 2 h exposure yielding the highest artemisinin content among other exposure hours but was marginally low compared to the untreated cultures. The treated cell suspension cultures showed no significant effect on the accumulation of artemisinin (Table 1). On the other hand, suspension cultures treated with varying DMSO concentrations did not show any enhancement in artemisinin production as well, in which case artemisinin is virtually absent (Table 2).

**Table 1.**
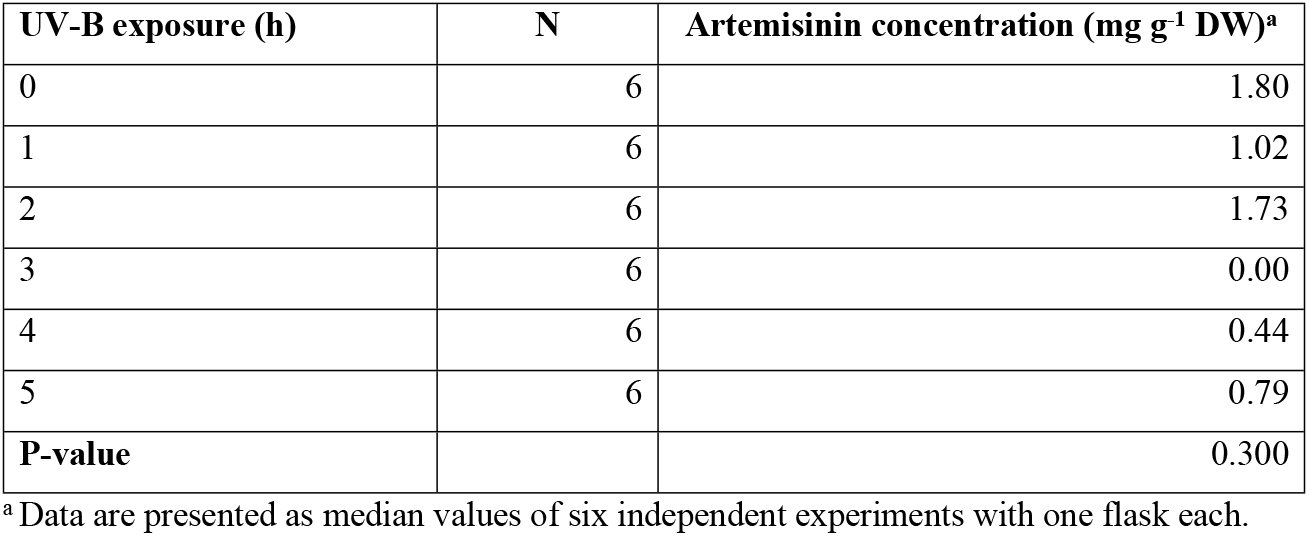
Artemisinin concentration in cell suspension cultures after the application of stress treatment with UV-B radiation at different durations.

| UV-B exposure (h) | N | Artemisinin concentration (mg g<sup>-1</sup> DW)<sup>a</sup> |
| --- | --- | --- |
| 0 | 6 | 1.80 |
| 1 | 6 | 1.02 |
| 2 | 6 | 1.73 |
| 3 | 6 | 0.00 |
| 4 | 6 | 0.44 |
| 5 | 6 | 0.79 |
| **P-value** |  | 0.300 |

<sup>a</sup> Data are presented as median values of six independent experiments with one flask each.

**Table 2.** Effect of different concentrations of DMSO elicitation on artemisinin accumulation.

| DMSO concentration (% [v/v]) | N | Artemisinin concentration (mg g<sup>-1</sup> DW)<sup>a</sup> |
| --- | --- | --- |
| 0.00 | 9 | 0.00 |
| 0.10 | 9 | 0.00 |
| 0.25 | 9 | 0.00 |
| 0.50 | 9 | 0.00 |
| 1.00 | 9 | 0.00 |
| 2.00 | 9 | 0.00 |
| **P-value** |  | 0.659 |

<sup>a</sup> Data are presented as median values of nine independent experiments with one flask each.

Viewed together, the hypothesis that these applied stress treatments would stimulate the enhancement of artemisinin production in cell cultures of a low producer of artemisinin is rejected. This lack of stimulation seemed to be tied to several physiological attributes that could possibly explain the roadblock responsible for failure. The physiological state of cultures in terms of age, the presence of glandular trichomes (a feature closely related to age factor) and the intrinsic regulation of cellular metabolic homeostasis were identified as important factors that may influence stress responses of *A. annua* that in turn results in a distortion of elicitation outcome. At present, our findings are in complete agreement with former experiments that suggested the importance of glandular secretory trichomes in the context of artemisinin production and sequestration. The suspension culture of *A. annua* cells lack cellular or tissue differentiation which consisted of only free cells and cell aggregates (Fig 2B). Spatial distribution of alkaloid biosynthesis is commonplace in whole plants, and with this secondary metabolite synthesis and accumulation are often correlated with cytodifferentiation. For example, the undifferentiated *Papaver somniferum* cell cultures failed to produce morphine, as it was noted that laticifers are absent from these cultures but present in seedling cultures [51–53]. Another related example of tissue-specific metabolite is the anti-depressant hypericin and hyperforin derived from *Hypericum perforatum*. The undifferentiated cells are unable to produce these phytochemicals without a certain extent of morphogenesis [54]. In the case of *A. annua*, whereby localization of enzymes specific to artemisinin biosynthesis in the apical cells of glandular secretory trichomes is evident, the differentiation and development of specialized cell types important for their defensive function in cell cultures are yet to occur unless otherwise induced to undergo *in vitro* differentiation [55–57]. However, this reasoning does not seem to apply to former elicitation experiments on *A. annua* suspension cultures. While Ferreira and Janick reported either trace or no artemisinin in HAP cell cultures (parallel to our observation) [58], Baldi and Dixit, Chan et al., Caretto et al. and Salehi et al. were able to stimulate artemisinin production by several folds via the incorporation of mevalonic acid lactone, methyl jasmonate, potassium nitrate and combination of coronatine and sorbitol [32–34, 59].

From observations of previous studies, it may be surmised that the limited effect of these stress-inducing agents is associated with feedback inhibition of product synthesis. The exertion of feedback control on artemisinin production was clearly demonstrated by Arsenault et al. on mature *A. annua* when exogenous artemisinin was applied, acting through the repression of transcription of cytochrome P450 (CYP), *CYP71AV1* and/or *ADS* genes [60]. As plant cells are constantly bathed in the liquid medium, it is likely that extracellular artemisinin that was excreted out of the cells into the medium, due to the lack of trichomes for compartmentalization, may trigger the negative feedback mechanism to inhibit excess production of this phytotoxic metabolite [61]. Similar observations were made in Soetaert et al. [62]. The limited effects of exogenous jasmonic acid on Anamed A3 cultivar (HAP chemotype) was reasoned to be a result of endogenous jasmonic acid-signaling reaching its maximum effect, thus impeding any additional response. It is also important to note that the low accumulation of artemisinin in cell suspension cultures can be affected by enzymatic or non-enzymatic degradation in the medium [63]. As the culture fluid functions as an extracellular lytic compartment due to the release of catabolic enzymes and acids from cells [64], the synthesized artemisinin will be rapidly metabolized as its specialized sequestration site (the subcuticular space of glandular secretory trichomes) is obviously missing in the present study. Such detoxification of excreted toxic alkaloid has been described by Weiss et al. in cell suspension cultures of *Eschscholzia californica* [65]. With the addition of sanguinarine to the suspension culture, a rapid decline and eventual disappearance of the benzophenanthridine alkaloid from the medium coincided with an increase of the less toxic compound, dihydrosanguinarine.

Given the difficulties in improving artemisinin accumulation of a LAP chemotype in a cell suspension system, we postulated that the failure of secondary metabolite elicitation was attributed to: (i) the general lack of cytodifferentiation of cell cultures; (ii) negative feedback regulation of artemisinin biosynthesis; and (iii) artemisinin sequestration by cellular detoxification. With these in mind, further extraction studies of artemisinin were performed on culture media post-treatment. The absence of the antimalarial compound in cell culture medium is indicative that artemisinin is not excreted into the medium by the producing cells, and it would also suggest that artemisinin biosynthesis did not occur in either elicited or non-elicited cells. This reaffirms the assumption that *A. annua* cells lacking specialized tissues are unable to synthesize and accumulate artemisinin as reflected by the negative results obtained from the DMSO elicitation study. Furthermore, assuming that the cultured cells are capable of artemisinin production and these products are released into the medium due to the lack of storage tissues, the prolonged incubation period after the addition of DMSO may have resulted in artemisinin degradation by hydrolytic and oxidizing enzymes present in the culture medium. Woerdenbag et al. reported of the rapid decomposition of artemisinin when incubated with a cell homogenate of *A. annua* or spent culture medium, and in a solution of commercial horseradish peroxidase as control [66]. The very low artemisinin content in undifferentiated cell cultures of *A. annua* could be negatively impacted by extracellular peroxidase activity in the surrounding medium. Clearly, more detailed analysis should be made regarding this aspect to elucidate the possibilities of enzymatic hydrolysis of artemisinin in cell culture systems, and thus providing opportunities for further improvement in minimizing the loss of artemisinin in such *in vitro* production system.

In the case of UV-B elicitation, however, artemisinin was detected in cell extracts (albeit at low levels and fluctuations) but remained undetected in the medium. These findings appear to support the assumption that an as-yet-unknown self-resistance mechanism towards self-produced phytotoxic metabolite may exist in *A. annua* cell cultures, at least for this European variety. The fluctuating cellular artemisinin contents seemed to parallel the recycling mechanism of sanguinarine detoxification. Presumably, excreted artemisinin may be rapidly reabsorbed by producing cells and converted into a less toxic product via reduction reactions, in which case the resulting compound may then undergo further biosynthetic reactions [65]. Although there is no direct evidence regarding self-resistance in *A. annua* suspension culture, but there remains substantial scope for further research to unravel the mechanism underlying protection against self-intoxication in cells (if any); and provide a deeper insight on the differential control of artemisinin sequestration in relation to developmental stage (cell cultures and intact plants). Of note, we believe that the regulatory mechanism underlying artemisinin production is closely linked to the self-resistance mechanism [67]. Thus, it would be worthwhile to warrant further studies to elucidate the relationship between artemisinin biosynthetic pathway and its detoxification process in cell cultures.

Lastly, we extended our examination to determine whether the degree of cytodifferentiation of cell cultures affects artemisinin biosynthesis genes. Conventional PCR assays revealed that 3 out of 4 artemisinin specific biosynthetic genes, ADS, *DBR2* and *ALDH1* were absent in cultured cells. To ascertain the underlying cause for gene absence, whether variation in chemotypic characters of this particular *A. annua* variety or the developmental state of the cultures could have influenced the regulation of artemisinin biosynthesis, soil-grown plants of the corresponding chemotype were used as a comparison and it was found that those genes were expressed in juvenile leaves (Fig 4). Thus, these observations suggested the importance of developmental timing on the spatial-temporal regulation of gene expression influencing artemisinin biosynthesis. Similar to previous description of cell suspension cultures of *Lupinus polyphyllus, Cytisus scoparius* and *Laburnam alpinum*, quinolizidine alkaloids were produced in orders of magnitude lower (0.01 – 5 μg g^-1^ fresh weight) compared to those of differentiated plants (500 – 5000 μg g^-1^ fresh weight). This was inferred that genes responsible for quinolizidine alkaloid formation were expressed at very low levels [68]. Likewise, the inability to detect *ADS* expression in *A. annua* cell cultures has also been reported by Caretto et al. and Jing et al. [34, 69], although unfortunately not all genes of the pathway were studied. Our results were consistent with these reports but were in disagreement with Wallaart et al. [70], who reported that *ADS* was not detected in young leaves of unstressed *A. annua* plants. Furthermore, as *ADS* is an essential gene for the first committed step of artemisinin formation [71] and the modest effect of yeast expressing only *ADS* on amorpha-4,11-diene production [72], its absence in cultured *A. annua* cells would likely lead to the truncation of artemisinin biosynthesis pathway. Lacking genes encoding both DBR2 and ALDH1 enzymes necessary for the preceding steps in the pathway would further disrupt artemisinin biogenesis. It is also noteworthy that our observations are consistent with previous data suggesting the deployment of divergent and distinctive classes of CYP/CPRs in *A. annua* plants to cope with the reductive demand of P450-catalyzed reactions [73]. Since CYP71AV1 is a membrane-bound, multifunctional enzyme that is involved in a wide range of oxidative metabolic reactions, such as demonstrated by its efficient *in vivo* conversion of amorpha-4,11-diene to artemisinic acid by recombinant CYP71AV1 [72], it is therefore no surprise that both CYP71AV1 and CPR (a redox partner of the former) genes are present even in cultured cells.

**Fig 4.**
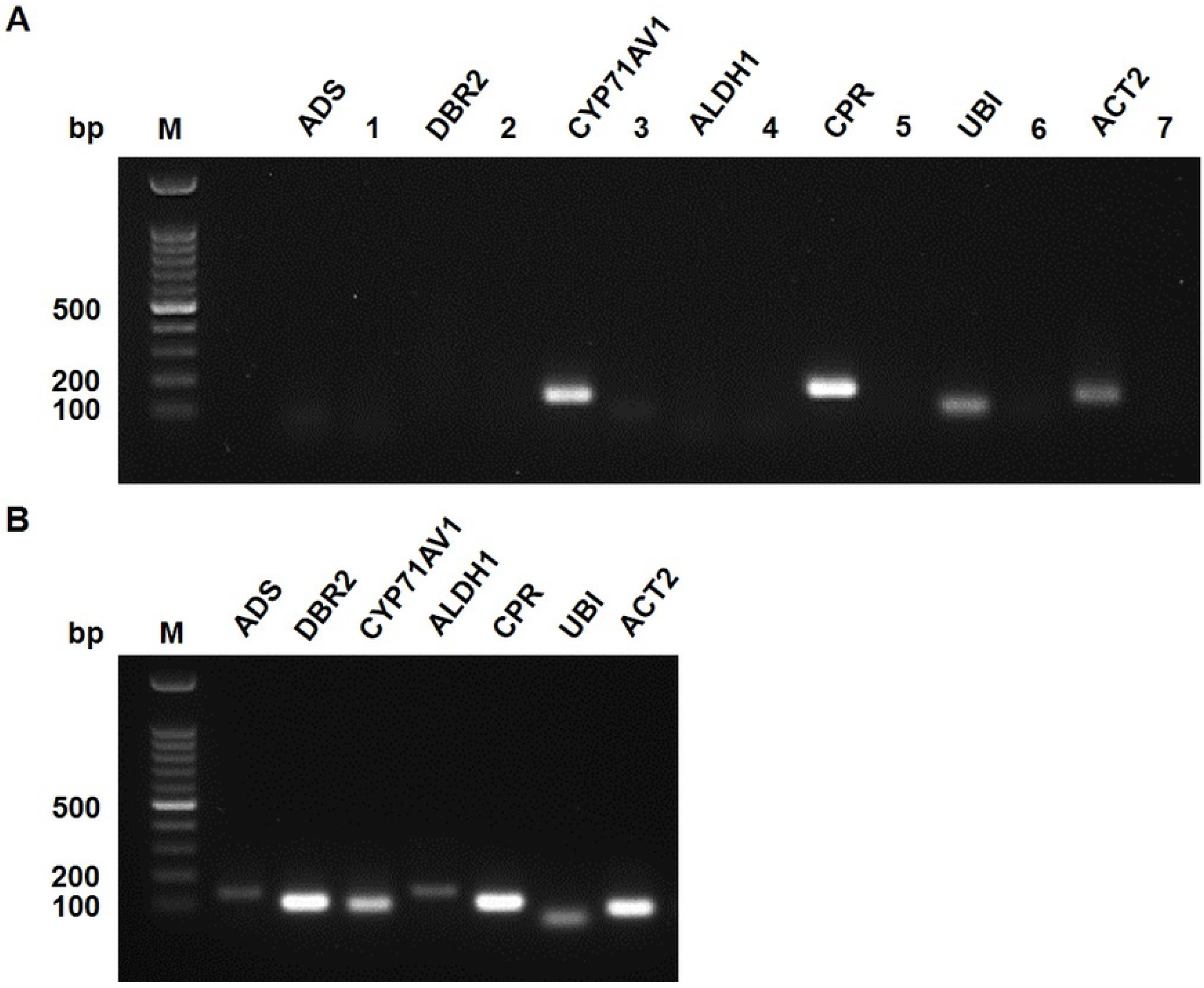
Comparison of PCR profiles of amplified artemisinin biosynthetic genes and housekeeping genes. (A) Untreated cultured cells and (B) leaf tissues from soil-grown *A. annua* using gene-specific primers. M, 100 bp DNA ladder; 1-7, no reverse transcriptase control of respective genes.

Taken together, it is reasonable to conclude that the inefficiency of elicitation in the present study was not due to the incompatibility or choice of stress-inducing agents, but rather the spatial-temporal regulation of artemisinin biosynthetic pathway which is associated with the lack of cellular differentiation in *A. annua* cell suspension cultures. Apart from the repression of most key genes along the pathway, other processes such as transportation, accumulation and degradation of artemisinin should be considered in determining whether a LAP plant-derived cell culture actually produces the antimalarial compound or not. Details of the metabolism of artemisinin in cell cultures, regardless of chemotype, remains a topic that deserve further investigation to evaluate these processes which in addition to biosynthesis are important for artemisinin production.

## Acknowledgements

The authors are grateful to Dr Ajit Singh for helpful advice on statistical analysis.

## Supporting information

**S1 Table. Preliminary data from elicitation trials on soil-grown LAP *A. annua* plants.**

**S2 Table. Gene-specific primer sequences.**

**S3 Table. Dataset of elicitation-induced plant cell production of artemisinin in liquid cell suspension cultures.**

